# orfipy: a fast and flexible tool for extracting ORFs

**DOI:** 10.1101/2020.10.20.348052

**Authors:** Urminder Singh, Eve Syrkin Wurtele

## Abstract

**Summary:** Searching for ORFs in transcripts is a critical step prior to annotating coding regions in newly-sequenced genomes and to search for alternative reading frames within known genes. With the tremendous increase in RNA-Seq data, faster tools are needed to handle large input datasets. These tools should be versatile enough to fine-tune search criteria and allow efficient downstream analysis. Here we present a new python based tool, `orfipy`, which allows the user to flexibly search for open reading frames in fasta sequences. The search is rapid and is fully customizable, with a choice of Fasta and BED output formats.

**Availability and implementation:** `orfipy` is implemented in python and is compatible with python v3.6 and higher. Source code: https://github.com/urmi-21/orfipy. Installation: from the source, or via PyPi (https://pypi.org/project/orfipy) or bioconda (https://anaconda.org/bioconda/orfipy).

**Supplementary information:** Supplementary data are available at https://github.com/urmi-21/orfipy

## Introduction

Open reading frames (ORFs) are sequences that have potential to be translated into proteins. They are delineated by start sites, at which translation is initiated by assembly of a ribosome complex, and stop sites, at which translation is terminated and the ribosome complex disassembles (1). Accurate annotation of the protein coding regions in a sequenced genome remains a challenging task in bioinformatics. Transcriptomic data is critical to address this challenge (2–4) Finding ORFs in transcripts is the first step to annotate the coding regions of genes.

Transcriptomic data can be exploited to identify previously unannotated genes, such as novel protein coding regions, alternative splicing events, and gene fusions (5). These data are also key to identifying orphan genes (2, 6), genes that are unique to a species without any homology to genes in other species (2, 6–9). Standard *ab initio* gene-prediction models trained on features of protein coding genes do not work well for identifying orphan genes, which are often shorter in length and sparse in canonical gene features (2, 10, 11).

Here we present `orfipy`, an efficient, intuitive tool for extracting ORFs from fasta files of sequence data. `orfipy` provides rapid, flexible search and multiple output formats to allow easy downstream analysis of ORFs. `orfipy` is developed for big transcriptomic datasets, in which annotating CDS is a common task. `orfipy` is particularly fast for multiple smaller sequences, such as transcriptomic and meta-genomic data.

## Implementation

`orfipy` is written in python, with the core ORF search algorithm implemented in cython to achieve faster execution times. `orfipy` efficiently searches for the start and stop codon positions in a sequence using the Aho–Corasick string-searching algorithm via the pyahocorasick library (https://pypi.org/project/pyahocorasick/). Vectorized implementation of ORF searching further improves the performance. `orfipy` can leverage multiple cpu-cores in order to process fasta sequences in parallel. The number of sequences to be processed in parallel is automatically estimated by `orfipy` at runtime, as based on available memory and cpu cores (See Supplementary Data).

### Input and output

`orfipy` takes nucleotide sequences in a multi-fasta file as input. Using the python package pyfaidx (13), `orfipy` creates an index from the input fasta file for easy and efficient access to the input sequences. In addition to the input sequences, users can provide input parameters that include minimum and maximum size of ORFs, list of start and stop codons, and/or a user-defined codon table (See Supplementary Data).

For easy, efficient and flexible downstream analysis, `orfipy` provides multiple output types. These are nucleotide or peptide sequences as fasta files, or coordinates of ORFs in BED format (Figure 1A). Because the number of input sequences can be very large when analyzing transcripts from meta-assemblies or large genomes, writing ORFs as fasta files can take take a lot of disk space; BED files reduce disk space use by storing only the coordinates of the ORFs. Moreover, using BED coordinate files can be helpful in developing more scalable and flexible downstream analysis pipelines. `orfipy` adds to the output file extra relevant information about codon use and ORF types; it can also group the output according to the longest ORF contained in each transcript or list each reading frame in each transcript (Figure 1A, B).

**Fig. 1.**
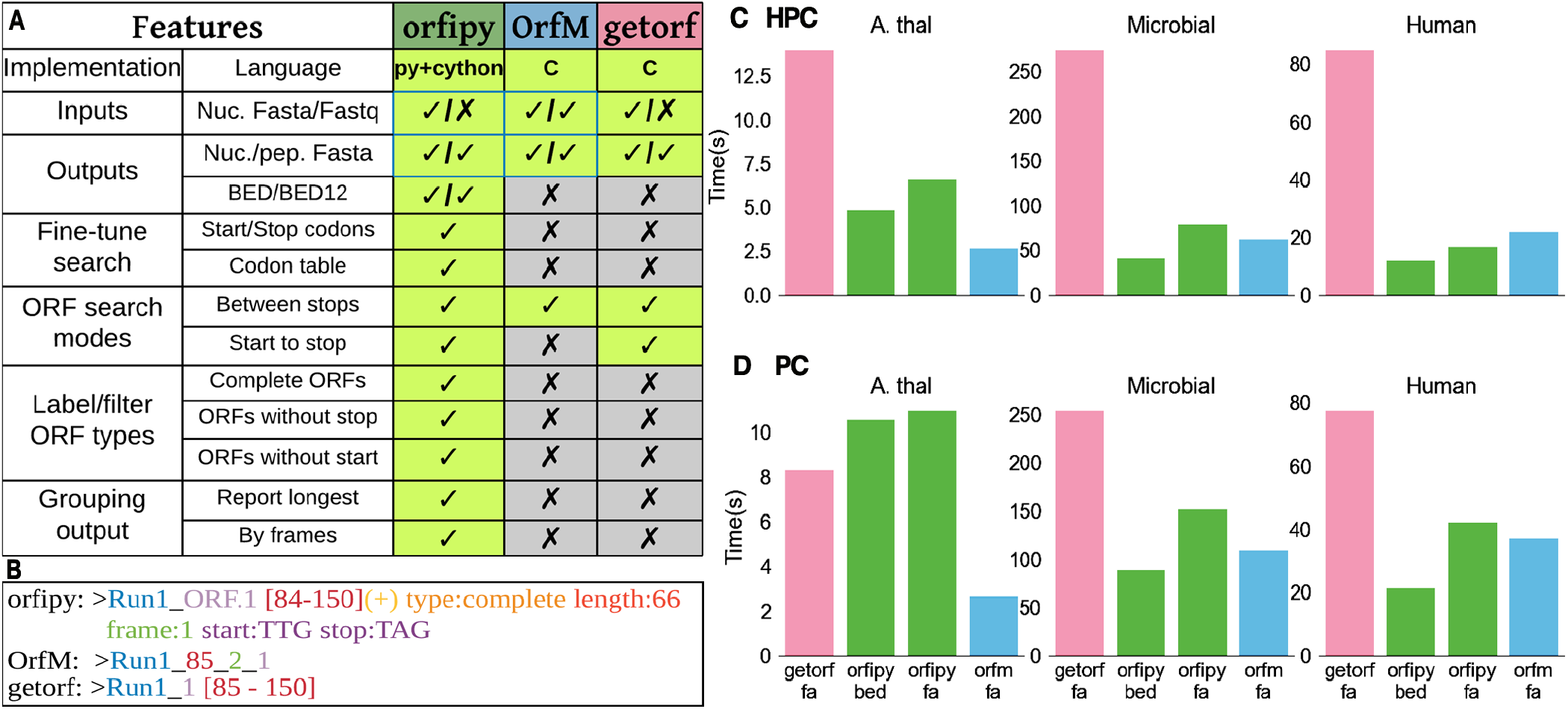
Comparison of orfipy features and performance with getorf and OrfM. We compared attributes of `orfipy` with two commonly-used tools for ORF identification. **A.**Comparison of `orfipy` features with getorf and OrfM. `orfipy` provides a number of options to fine-tune ORF search, this includes labeling the ORF type, reporting only the longest ORF, and reporting ORFs by translation frame. **B.**Example of fasta headers written to output files by each tool. `orfipy` output provides information about each ORF that can be readily used in downstream analyses. **C and D.**Runtimes on HPC (128 GB RAM; 28 cores) (**C**) and PC (8 GB RAM; 4 cores) (**D**) environments. Each analysis was run three times, via pyrpipe (12), and the mean runtime is reported. `orfipy` runtimes are faster than or comparable to `OrfM` for the large microbial and human transcriptome data. An HPC environment, with more cpu cores and RAM, decreased runtimes for `orfipy` and `OrfM` . `orfipy` is fastest when ORFs are saved to a BED file. Data sizes: *A. thaliana* genome 120 MB; microbial sequences 1.5 GB; human transcriptome 370 MB. fa: output ORFs as nucleotide and peptide fasta; bed: output ORFs as BED.

### Flexible ORF search

`orfipy` enables researchers to fully fine-tune ORF searches using a variety of options (Figure 1A). For example, users can limit ORF searching to a specific start codon or choose to output ORFs without an inframe start codon. `orfipy` labels each ORF as complete, 5-prime partial (lacking a start codon), or 3-prime partial (lacking a stop codon) for users to easily comprehend the results (See Supplementary Data).

### Comparison with existing tools

We compared `orfipy` with the popular tools getorf (14) and OrfM (15) (Figure 1). What sets `orfipy` apart from existing tools is its flexibility and the various options to fine-tune ORF searches and output (Figure 1A, B). We measured the runtimes to find and save ORFS in three datasets of different sizes and structures (Figure 1C,D) (See Supplementary Data). For a side-by-side comparison of each tool, we timed outputs to the nucleotide and peptide sequence format; we also timed outputs in the versatile BED format for `orfipy` (the other tools do not provide this option).

The runtime for each tool depends on the tool, the environment, the output type selected, and the size and structure of the data analyzed. In all scenarios except using a PC to analyse the smaller *A. thaliana* dataset, `orfipy` is much faster than `getorf`, and faster than or similar to that of `OrfM` . For the two larger datasets, runtimes are fastest using `orfipy` with BED output.

## Funding

This work is funded in part by National Science Foundation grant IOS 1546858, Orphan Genes: An Untapped Genetic Reservoir of Novel Traits, and by the Center for Metabolic Biology, Iowa State University.

## Notes

https://github.com/urmi-21/orfipy

### Competing Interest Statement

The authors have declared no competing interest.

## Bibliography

1. Shea J Andrews and Joseph A Rothnagel. Emerging evidence for functional peptides encoded by short open reading frames. Nature Reviews Genetics, 15(3):193–204, 2014.

2. Arun S Seetharam, Urminder Singh, Jing Li, Priyanka Bhandary, Zebulun Arendsee, and Eve Syrkin Wurtele. Maximizing prediction of orphan genes in assembled genomes. BioRxiv, 2019.

3. Thomas F Martinez, Qian Chu, Cynthia Donaldson, Dan Tan, Maxim N Shokhirev, and Alan Saghatelian. Accurate annotation of human protein-coding small open reading frames. Nature chemical biology, 16(4):458–468, 2020.

4. Khalid Mahmood, Jihad Orabi, Peter Skov Kristensen, Pernille Sarup, Lise Nistrup Jørgensen, and Ahmed Jahoor. De novo transcriptome assembly, functional annotation, and expression profiling of rye (secale cereale l.) hybrids inoculated with ergot (claviceps purpurea). Scientific reports, 10(1):1–16, 2020.

5. Zhong Wang, Mark Gerstein, and Michael Snyder. Rna-seq: a revolutionary tool for transcriptomics. Nature reviews genetics, 10(1):57, 2009.

6. Jing Li, Urminder Singh, Zebulun Arendsee, and Eve Syrkin Wurtele. Landscape of the dark transcriptome revealed through re-mining massive rna-seq data. bioRxiv, page 671263, 2020.

7. Diethard Tautz and Tomislav Domazet-Lošo. The evolutionary origin of orphan genes. Nature Reviews Genetics, 12(10):692–702, 2011.

8. Urminder Singh and Eve Syrkin Wurtele. Genetic novelty: How new genes are born. Elife, 9:e55136, 2020.

9. Nikolaos Vakirlis, Anne-Ruxandra Carvunis, and Aoife McLysaght. Synteny-based analyses indicate that sequence divergence is not the main source of orphan genes. Elife, 9:e53500, 2020.

10. Jorge Ruiz-Orera, Jessica Hernandez-Rodriguez, Cristina Chiva, Eduard Sabidó, Ivanela Kondova, Ronald Bontrop, Tomàs Marqués-Bonet, and M Mar Albà. Origins of de novo genes in human and chimpanzee. PLoS genetics, 11(12):e1005721, 2015.

11. Brennen Heames, Jonathan Schmitz, and Erich Bornberg-Bauer. A continuum of evolving de novo genes drives protein-coding novelty in drosophila. Journal of molecular evolution, pages 1–17, 2020.

12. Urminder Singh, Jing Li, Arun Seetharam, and Eve Syrkin Wurtele. pyrpipe: a python package for rna-seq workflows. bioRxiv, 2020. doi: 10.1101/2020.03.04.925818.

13. Matthew D Shirley, Zhaorong Ma, Brent S Pedersen, and Sarah J Wheelan. Efficient“ pythonic” access to fasta files using pyfaidx. Technical report, PeerJ PrePrints, 2015.

14. Peter Rice, Ian Longden, and Alan Bleasby. Emboss: the european molecular biology open software suite, 2000.

15. Ben J. Woodcroft, Joel A. Boyd, and Gene W. Tyson. OrfM: a fast open reading frame predictor for metagenomic data. Bioinformatics, 32(17):2702–2703, 05 2016. ISSN 1367-4803. doi: 10.1093/bioinformatics/btw241.

