## Supplementary Data for "orfipy: a fast and flexible tool for extracting ORFs"

### Supplementary Information

#### Getting started

Current stable version of `orfipy` can be installed via PyPI using the following command

```
pip install orfipy
```

Information on installing development version and source code is available from https://github.com/urmi-21/orfipy. During submission of this manuscript, the source code, can also be accessed here: http://doi.org/10.5281/zenodo.4091773.

#### Basic usage

`orfipy` requires a fasta file as input to run. Other several options can be used fine tune ORF search. Basic command to execute `orfipy` looks like:

```
orfipy [<options>] <in.fa>
```

`orfipy` help menu, which describes all the input options, can be printed using the command

```
orfipy -h
```

#### Providing start, stop codons and translation table

With `orfipy`, users can specify one of the 23 codon tables from NCBI (https://www.ncbi.nlm.nih.gov/Taxonomy/Utils/wprintgc.cgi?chapter=cgencodes) to use for translation. If needed, users can also input their own table with the –table parameter. The codon table should be in a .json file to be read by orfipy. An example is given here: https://raw.githubusercontent.com/urmi-21/orfipy/master/scripts/example_user_table.json
By default, the codon tables specify the start and stop codons to use. User can easily override these values using −−*start* and −−*stop* parameters when running orfipy. For example:

```
orfipy in.fasta --start TTG,CTG #uses TTG and CTG start codons
or
orfipy in.fasta --start ATG --stop TAA #uses only TAA as stop codon
```

### ORF searching modes

orfipy has two ORF searching modes to allow user to search for ORFs defined as:

1. Between Start and Stop: These ORFs are defined as the regions between a start and a stop codon. For example, consider the sequence

   AAAATGTTTAAAGGGCCCTAGTTT

   ORF starting with the start codon "ATG" and ending with the stop codon "TAG" is (stop codon not included in the ORF):

   ATG TTT AAA GGG CCC

2. Between Stops (regions free of stop codon): These ORFs are defined as regions free of stop codons. For examples, for the previous sequence, the ORFs reported in frame +1 will now be:

   AAA ATG TTT AAA GGG CCC
   and
   TTT

### ORF types

orfipy labels the ORFs as following types:

1. complete: This type of ORF has both a start and a stop codon. For example (stop codon included in ORF for reference):

   ATG TTT AAA GGG CCC TAG (STOP)

2. 5-prime partial: This type of ORF has a stop codon but lacks a start codon. For example:

   TTT AAA GGG CCC TAG (STOP)

3. 3-prime partial: This type of ORF has a start codon but lacks a stop codon. For example:

   ATG TTT AAA GGG CCC

By default orfipy outputs all the complete ORFs. Users can use the flags $--partial-5$ and $--partial-3$ to include the 5-prime partial and 3-prime partial ORFs.

### Memory considerations

orfipy loads multiple fasta sequences into memory and searches for ORFs in parallel. This approach is particularly faster for files containing multiple smaller sequences such as de-novo transcriptome assembly data. For files with much larger sequences such as whole genomes, the amount of memory required by orfipy increases due to the large sequence size and number of ORFs found (which are proportional to sequence size). In such scenarios orfipy can use a lot of memory and may affect system's performance. To deal with this user can set the $--chunk-size$ option to have control over the memory usage. orfipy will process the input file in chunks of size $--chunk-size$. By default orfipy is able to determine an optimal value of $--chunk-size$ based of system's available memory, cpu cores and the input file provided.

### Benchmarking details

To compare the runtimes of `orfipy` with getorf[1] and OrfM[2], we used three datasets: *A. thaliana* whole genome (TAIR 10), 5,000 microbial genomes and RNA, and Human transcriptome (Ensembl release-101). Each tool was run to find ORFs defined as regions free of stop codons as OrfM only supports this mode. Each tool was run to save ORFs as nucleotide and peptide fasta files. Additionally, `orfipy` was also run to output ORFs in BED format. For *A. thaliana* minimum ORF size was set as 300 while for other two datasets minimum ORF size was set to 99. A script was written to execute the tools were executed via pyrpipe[3] to capture the runtimes. Tools were run three times and mean runtime for each tool is reported. Code to run the benchmarks is available at https://github.com/urmi-21/orfipy/tree/master/scripts.

Basic commands for executing tools looked like the following:

**orfipy**

```
orfipy --min <m> --dna <nuc.fa> --pep <pep.fa> --between-stops <in.fa>
and
orfipy --min <m> --bed <out.bed> --between-stops <in.fa>
```

**OrfM**

```
orfm -m <m> -t <nuc.fa> <in.fa> > <pep.fa>
```

**getorf**

```
getorf -find 0 -min <m> -outseq <pep.fa> -sequence <in.fa>
getorf -find 2 -min <m> -outseq <nuc.fa> -sequence <in.fa>
```

### Compatibility with other tools

This section describes how to set `orfipy` parameters to extract the same ORFs as with some existing tools. This will be helpful to users wanting to switch to orfipy from existing tools.

#### *OrfM*

OrfM[2] has only one run mode i.e. find ORFs defined as free of stop codons. To use `orfipy` to extract these ORFs, use the flag $--between-stops$. OrfM command:

```
orfm -m <m> -t <nuc.fa> <in.fa> > <pep.fa>
```

`orfipy` command:

```
orfipy --min <m> --dna <nuc.fa> --pep <pep.fa> --between-stops <in.fa>
```

#### *getorf*

getorf[1] can find ORFs defined as regions free of stop codons or regions between start and stop codons. User can choose among these options by using the $-find$ parameters in getorf. When $-find$ is set to 0 or 2 (0 for peptide output and 2 for nucleotide), getorf extracts ORFs defined as regions free of stop codons. This is same as OrfM and `orfipy` command will be identical to what described for OrfM i.e. using the $--between-stops$ flag.

When $-find$ is set to 1 or 3 (1 for peptide output and 3 for nucleotide), getorf extracts ORFs defined as regions between start and stop codons. However, getorf only uses $ATG$ as the start codon asn also outputs ORFs lacking a stop codon. Thus, to get same ORFs as getorf from `orfipy`, users can specify the start codon to be ATG and to include 3-prime partial ORFs (ORFs lacking a stop codon).

getorf:

```
getorf -find 3 -min $3 -outseq "$2/getorf_3_d" -sequence $1
```

Equivalent `orfipy` command will be:

```
orfipy --min <m> --dna <nuc.fa> --start ATG --partial-3 <in.fa>
```

#### *TransDecoder*

The TransDecoder[4] tool can be used for identification of candidate coding sequences. TransDecoder.LongOrfs command identifies long ORFs in sequences and then computes a score for each ORF. Transdecoder.Predict then predicts the likely coding regions using score and length based filtering criteria. Transdecoder.Predict outputs ORFs codrdinate in BED12 format. `orfipy` can out output ORFs in the same format, by choosing out type as BED12 and using $--include-stop$ flag.

```
orfipy --bed12 <orfs.bed> --start ATG --partial-3 --partial-5 --include-stop <in.fa>
```

**Note:** Since TransDecoder applies a score based criteria, the number of ORFs reported by TransDecoder.Predict will be different than `orfipy`.

### References


1. Rice, P., Longden, I. & Bleasby, A. Emboss: the european molecular biology open software suite (2000).
2. Woodcroft, B. J., Boyd, J. A. & Tyson, G. W. OrfM: a fast open reading frame predictor for metagenomic data. *Bioinformatics* **32**, 2702–2703, DOI: 10.1093/bioinformatics/btw241 (2016). https://academic.oup.com/bioinformatics/article-pdf/32/17/2702/17345985/btw241.pdf.
3. Singh, U., Li, J., Seetharam, A. & Wurtele, E. S. pyrpipe: a python package for rna-seq workflows. *bioRxiv* DOI: 10.1101/2020.03.04.925818 (2020). https://www.biorxiv.org/content/early/2020/04/08/2020.03.04.925818.full.pdf.
4. Haas, B. & Papanicolaou, A. Transdecoder (2017).
